# When Stopping Requires Going; Physiological Similarities Between Action Cancellation and the Cancellation of Action Cancellation

**DOI:** 10.1101/2025.01.07.631825

**Authors:** Simon Weber, Sauro E. Salomoni, Mark R. Hinder

**Author notes:** Corresponding author: Simon Weber.

## Abstract

The reactive cancellation of real-world actions typically requires complex combinations of both muscle contraction and/or muscle suppression. However, current experimental paradigms solely examine contexts in which action cancellation requires muscle suppression. To provide fundamental insights into inhibitory control mechanisms, we directly compared the latency of action cancellation in novel paradigms where ‘stopping’ required either suppression of planned activation or reinstatement of ongoing activity.

20 healthy adults (mean age = 32.2 years) completed novel variants of the stop signal task (SST) in which each trial began with tonic force production to depress two buttons. When a go signal appeared, participants were required to release these buttons. On a subset of trials, a stop signal occurred after a brief delay, and participants were required to cancel the release of one of the buttons. Data in these variants were compared to conventional response-selective SSTs, in which the go signal required bilateral button presses and stop signals necessitated cancellation of one of these responses. Electromyographic (EMG) recordings allowed detailed comparison of the characteristics of muscle contraction and suppression (i.e., stopping speed) across these tasks.

When physiological evidence of synchronous action cancellation in both hands was observed (supporting recent models of complex stopping), EMG measures of action cancellation speed did not differ (p = 0.863, BF01 = 8.49) between cancellation of releases and cancellation of presses conditions. This result suggests that response inhibition may broadly characterise reactive control to maintain a current physiological state rather than specific cancellation of a voluntary response.

## 1. Introduction

### 1.1 Cancelling actions requires both suppression and activation of specific muscles

Action cancellation describes the neural suppression of planned or initiated movements in response to updated sensory information or decision outcomes. Everyday examples of action cancellation include suddenly halting a half-taken step when a cyclist crosses your path or retracting your hand when your notice an insect on an object you were reaching for. Notably, these scenarios require a combination of both muscle suppression (of the agonist muscles facilitating the initial movement) and activation (of opposing antagonist muscles; to actively counteract the momentum of the initial action). In the laboratory, stop signal tasks (SSTs) represent the gold-standard experimental paradigm for investigating action cancellation (Verbruggen et al., 2019). A typical SST involves a series of trials, the majority of which require participants to rapidly enact a prescribed movement (usually a simple button press with a finger) in response to a visual or auditory stimulus (the “go signal”). On a subset of trials, the go signal is followed by a “stop signal”, communicating that the participant’s prepotent response must be cancelled, such that they do not press the button. When cancelling an initiated finger press, healthy young adults typically begin to show a decrease in effector muscle activity (as recorded by electromyography; EMG) approximately 160ms following the stop signal (Jana et al., 2020; Raud & Huster, 2017, Raud et al., 2022; Salomoni et al., 2024). This empirical, single trial measure of stopping speed (usually referred to in the literature as “CancelTime”) is thought to reflect the endpoint of a neural cascade which underpins the cancellation of action. However, the precise neural correlates of this process are incompletely understood, and may vary depending on context.

Brain stimulation studies support the idea that action cancellation is a specialised motor process unique from the voluntary initiation of action. For instance, reactive stopping increases intracortical inhibition in the primary motor cortex (e.g., Coxon, et al., 2006; van den Wildenberg et al., 2010) and task-irrelevant muscles are supressed during reactive action cancellation, suggesting that a unique mechanism is at play (Badry et al., 2009; Greenhouse, et al., 2011; Majid, et al., 2012). Furthermore, unlike movement initiation, action cancellation may feature involuntary components that suppress motor output irrespective of whether or not this suppression is congruent with the intended response (Wessel & Aron, 2017). Specifically, our previous work showed that when an unexpected stimulus required enacting an additional action, rather than cancelling an action, muscle suppression was still observed (Weber et al., 2023). However, given the scenarios described above require a combination of activation *and* suppression of different muscle groups, this raises questions regarding whether inherent inhibitory mechanisms act as an impediment in contexts where cancelling an initial action predominantly requires muscle activation. To investigate this, we developed a novel version of the SST in which the go response required the *release* of tonic muscle activity and the cancellation of this response (following presentation of a stop signal) required an *increase* of muscle activity, to reestablish the initial level of tonic activation.

### 1.2. “global” or “selective” action cancellation

Global stopping (cancelling all ongoing movements) and response-selective stopping (the cancellation of only a subset of ongoing movements) may rely on distinct neural underpinnings. The latter is often considered to be more generalisable to reactive stopping in real-world scenarios, when the requisite action updating is complex (for instance cancelling a planned step while continuing the arm movements to bounce a basketball). One method for assessing the selectivity of inhibitory responses is to use a particular variety of SST, referred to in the literature as response-selective or motor-selective SSTs (Bissett, 2021). Stop trials in these tasks require participants to inhibit one component of a multi-component movement, while continuing with others. This typically involves cancelling one part of a bimanual response and completing a unimanual response on the non-cancelled side. Numerous theoretical accounts suggest that stopping all effectors (“global stopping”) relies on distinct neural correlates to response-selective stopping. For instance, it has been argued that a rapid hyperdirect basal ganglia pathway mediates global stopping, while the indirect pathway mediates response-selective stopping (Aron, 2011). Another recent framework suggests that an involuntary inhibitory response (referred to as a “pause”) occurs globally in response to attentional capture, while selective action cancellation can only be applied by a distinct, slower, and consciously controlled mechanism (referred to as a “cancel”; Diesburg & Wessel, 2021; Schmidt & Berke, 2017). However, recent computational modelling of behavioural data from response-selective stopping tasks is consistent with a global stop mechanism followed by the rapid initiation of a distinct action (for the component not cued to stop), rather than the implementation of a selective inhibition mechanism (Gronau, 2023). Here, we used response-selective stop signal tasks in order to investigate whether action cancellation (of either muscle activation or suppression) demonstrated evidence of global or selective inhibitory processes.

### 1.3. Voluntary stopping and attentional capture both suppress motor output

Growing evidence suggests two distinct inhibitory processes are triggered by the presentation of a stop signal. One describes the consciously controlled cancellation of movement and can be deliberately and selectively applied to only relevant effectors. The other is an involuntary inhibitory response that is not unique to action cancellation, and transiently suppresses all motor output following attentional capture to unexpected stimuli (Wessel & Aron, 2017 Sebastian et al., 2021). This transient interruption to motor output occurs ubiquitously in everyday scenarios. For instance, the rhythm of our speech may be interrupted mid-sentence because a pigeon unexpectedly flies into view. This transient pause to motor output occurs before we have consciously recognised that the unexpected stimulus is a pigeon, that it is irrelevant to our current task goals and thus should be ignored. Because stop signals involve both the voluntary cancellation of an action based on task goals and attentional capture to an unexpected stimulus these two sources of inhibition become conflated (Weber et al., 2023).

One experimental approach to distinguishing between these processes is to include additional infrequent signals, usually referred to as “ignore signals”, in an SST. Like the stop signals, these occur on a subset of trials and follow the go signal by a variable delay. However, *unlike* the stop signals they do not require the cancellation of movement. Instead, they require participants to continue with their initial response and ignore the additional stimulus. It can then be argued that these stimuli will only recruit the inhibitory processes associated with attentional capture, and not the voluntary, consciously controlled cancellation of movement. SSTs featuring stop and ignore signals are usually referred to as “stimulus selective” SSTs. Past research has observed that suppression of motor output is observable in response to ignore stimuli (Ko and Miller, 2013; Wadsley 2023a; Weber 2024). Furthermore, brain stimulation research has demonstrated that neural suppression of the corticospinal tract occurs similarly for stop and ignore signals until ∼200ms following the presentation of the stop signal, at which point suppression is released in ignore trials (Tatz, 2021). In the current experiment, ignore stimuli required the release of tonic contraction in one condition (the release-ignore condition, explained below), and an increase in muscle activity in another (the press-ignore condition). If the involuntary pause or attentional capture component of inhibition is involuntarily recruited in these trials, and this invariably results in muscle *suppression*, then response times in ignore trials requiring a release should be faster than those requiring a press (because the involuntary process is congruent, rather than counter, to the required action).

In sum, the current experiment sought to investigate whether involuntary inhibitory mechanisms act as an impediment in contexts where cancelling an initiated action predominantly requires muscle contraction, or whether stopping mechanisms can adapt to optimise action cancellation even when this involves muscle activation. This was investigated using response-selective and stimulus-selective stopping tasks.

## 2. Methods

### 2.1. Participants

20 healthy adults took part in the study (mean age = 32.2 years, SD = 8.6 years, range = 24-52; 8 male, 12 female). The number of participants was determined based on our previous research that was sufficiently powered to reveal statistically significant differences in CancelTime despite small effect sizes (Weber et al, 2023). Participants were recruited via the University of Tasmania’s School of Psychological Sciences research participation system and received either two hours of course credit (if required) or a $20 shopping voucher. All participants had normal or corrected to normal vision, and no known neuromuscular or neurological conditions or impairments. All participants provided informed consent. This research was approved by the Tasmanian Human Research Ethics Committee.

### 2.2. Computer and response interfaces

Participants were seated approximately 0.8 meters from a 27-inch, 240Hz computer monitor with forearms and hands pronated and resting on a desk, shoulder width apart. Each index finger rested against one of two custom-made response buttons. The buttons were mounted vertically, such that registering a response required participants to abduct their index fingers (inwards, on a horizontal plane), with the first dorsal interosseus (FDI) muscle on each hand acting as a primary agonist. This maximised FDI activation during voluntary button presses, increasing the signal-noise ratio in electromyographic data and facilitating the analysis of muscle activity. Button presses were registered via The Black Box Toolkit USB response pad and recorded using PsychoPy3 (Peirce et al., 2019).

### 2.3. Electromyography (EMG)

EMG recordings were made using disposable adhesive electrodes attached to the first dorsal interosseus (FDI) of participants’ left and right index fingers. Two electrodes were placed in a belly-tendon montage on each hand, with a third ground electrode placed on the head of the ulna. The analogue EMG signals were band-pass filtered at 20-1000 Hz, amplified 1000 times, and sampled at 2000Hz using Signal data acquisition board and amplifiers (Cambridge Electronic Design Ltd.). During the experiment, if recordings became noisy with background activity, the researcher would ask the participant to relax their hands between the voluntary responses.

### 2.4. Behavioural tasks overview

The experiment, developed and run using Psychopy3 (Peirce et al., 2019), featured four main conditions conducted in a single session of approximately two hours duration. The four conditions (described in detail below) were different variants of the stop signal task, differing in terms of the behavioural imperatives associated with the stimuli (i.e., whether the primary response required simultaneous button *presses*, or simultaneous button *releases*), and in terms of whether or not stimulus-selective stopping was required (i.e., whether or not ignore stimuli were included). The two conditions *without* stimulus-selective stopping are hereafter referred to as the “press condition”, and the “release condition”, while the two tasks requiring stimulus-selective stopping are referred to as the “press-ignore condition” and the “release-ignore condition” (on account of these tasks including “ignore trials”; described below).

The order of the four conditions was counterbalanced across participants to minimise practice and possible fatigue effects. The same pre-randomised trial order was used for the press and release conditions, to minimise any potential order effects from impacting comparison. A different sequence (that included ignore trials) was used for both the press-ignore and the release-ignore conditions. Trial order within each block was pre-randomised, with the criteria that no more than two infrequent trial types (stop and ignore trials, see below) occurred consecutively and the first trial immediately after a break was always a go trial. Trials (in all tasks) were separated by a 1 second inter-trial interval, and a rest break occurred after every 50 trials. During the breaks, text appeared asking participants to rest their hands and press one of the buttons once they were ready to continue. The break screen also displayed feedback indicating the proportion of correct responses in stop trials and average reaction time (RT) of correct go responses from the previous 50-trial block.

### 2.5. Go-only blocks

Prior to completing the main experimental conditions, participants completed two short “go-only” blocks, each consisting of 20 “go trials”. One of these blocks featured go trials requiring button presses (the “press go-only block”), while the other required button releases (the “release go-only block”). The go trials in the press go-only block began with a fixation cross appearing in the centre of the screen for a pre-randomised duration drawn from a truncated exponential distribution ranging from 250-1000ms^1^. Following this, the go stimulus would appear (two yellow circles, 3cm in diameter and 15 cm apart, equidistant from the centre of the screen). The imperative associated with the appearance of the go signal was for participants to press both buttons simultaneously. The response window (from when the go stimulus appeared, to when a response for that trial could no longer be recorded) lasted 1200ms.

In the release go-only block, the go trials required button releases. These trials began with a screen reading “hold both buttons”. This text remained on the screen until both buttons were depressed, and then (as with the trials described immediately above) a fixation cross appeared for 250-1000ms. The same go stimulus (two yellow) circles was then presented, though in this instance this necessitated participants’ *release* of both buttons simultaneously.

In both go-only blocks, and in go trials across all the main experimental conditions described below, the response period (always 1200ms) was immediately followed by a feedback screen displaying the reaction times for the left and right button responses for the duration of 1 second, before the 1 second inter-trial interval (blank black screen) occurred, followed by commencement of the next trial. The order in which participants completed the two go-only blocks was counterbalanced.

### 2.6. Press and press-ignore conditions

The press condition featured 100 go trials and 50 response-selective stop trials (described below and shown in Figure 1) occurring in a pre-randomised order. In the press condition, the go trials were the same as the go trials in the press go-only block. The stop trials were evenly distributed as left stops and right stops (25 of each). During stop trials the go signal appeared, though after a variable “stop signal delay” (SSD; explained below) one of the yellow circles was replaced by a “stop signal”. The stop signal was a blue or magenta circle (this was counterbalanced) appearing on either the left or right side of the screen and remained on screen until the end of the response period. This indicated that participants were required to inhibit their response on that side, while continuing their response on the other side (that is, response-selective stopping was required). If participants succeeded in selectively cancelling their movement (e.g., pressed only the left button when the stop signal occurred on the right), text appeared during the feedback window reading “correct”. If participants made any other response (pressed both buttons, failed to press either button, pressed the button on the wrong side) performance was deemed unsuccessful, and the feedback text displayed “incorrect”. Reaction times were not displayed in stop trials.

**Figure 1:**
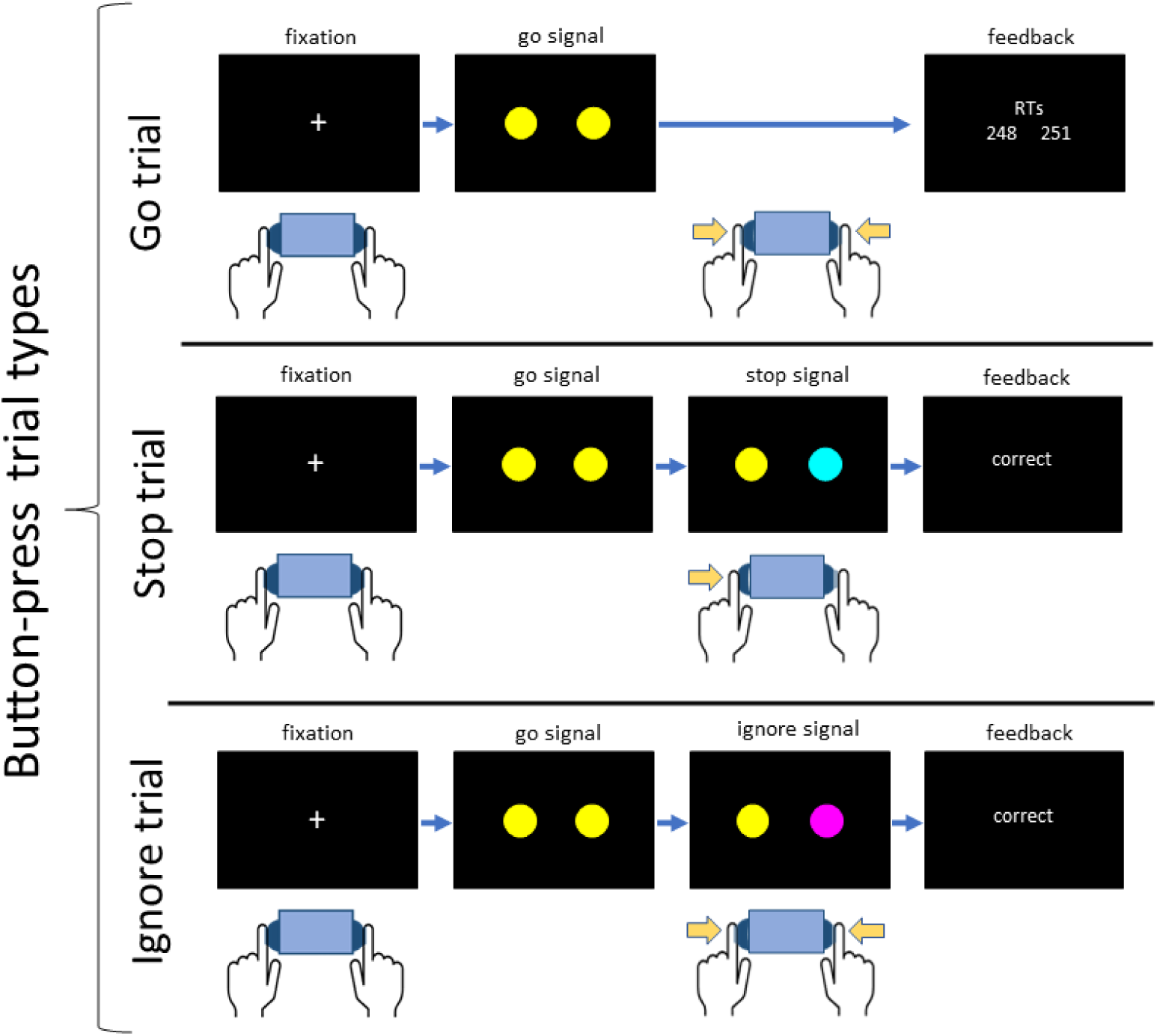
Trial types occurring in the two conditions requiring button presses as the primary response. Note that the “press” condition featured only go and stop trials, while the “press-ignore” condition also included ignore trials. Trials began with a fixation cross, while participants rested their fingers by the response buttons. The imperative associated with the go signal was to press both buttons simultaneously (button presses are indicated with arrows displayed next to the hand icons). The stop signal (here, a cyan circle) could appear on either the left or right side and occurred after a variable stop signal delay (SSD, described in main text). Participants were required to refrain from pressing the button on the side of the stop signal. Ignore signals (occurring in ignore trials, in the third row; magenta circle), also occurred after SSD, but required a bimanual button press, as in go trials.

The press-ignore condition featured 300 trials, of which 200 were go trials, 50 were stop trials and 50 were “ignore trials”. Go trials and stop trials followed the same procedure as those occurring in the press condition. As with the stop trials, ignore trials involved one of the yellow circles comprising the go stimulus changing colour following SSD. For half of the participants the colour used for ignore signals was magenta and for half it was cyan (whichever colour was not assigned to the stop trials for that participant was used for the ignore trials). During these trials, participants were required to ignore the colour change and continue with the button press (i.e., respond bimanually, as though both circles had remained yellow). Ignore trials, like stop trials, were evenly distributed as 25 left and 25 right ignores.

SSD refers to the time between the presentation of the go signal and the stop (or ignore) signal. Separate SSDs were used for left stop and right stop trials. Initially, both SSDs were set to 200ms, and this was incremented (shorter or longer; 50 ms increments) based on success in the previous stop trial of that condition. For example, following a correct right stop trial, the SSD for the subsequent right stop would increase by 50ms making stopping in the next stop trial more difficult. Following a failed right stop trial, the SSD would *decrease* by 50ms, making a subsequent right stop trial easier. Using this staircase procedure, the probability of successfully stopping should be approximately 50%, enabling assessment of both successful and failed stop trials. The timing of the ignore signals used the SSD that was set following the previous stop trial occurring on the same side (i.e., the timing of an ignore signal for a left ignore trial was the same as the SSD set following the last left stop trial). Performance on ignore trials did not influence SSDs.

In go trials, if participants responded more than 200ms slower than their average response time in the press go-only block, a message appeared during the feedback screen, above the RTs, reading “You’ve slowed down”. This feedback was included to minimise the degree to which participants would slow their responses in anticipation of stop signals (i.e., proactive slowing), a behavioural adjustment which can interfere with the SSD calculations described above, resulting in very high stop trial accuracy rates and invalidating SSRT calculations (Verbruggen et al., 2019).

### 2.7 Release and release-ignore conditions

The release condition featured 100 go trials and 50 stop trials occurring in a pre-randomised order. The go trials were the same as the go trials in the release go-only block (described above). In stop trials, the stop *stimulus* was the same as that used in the press condition (i.e., the same visual stimulus), but the imperative associated with that stimulus differed. Specifically, when the stop signal appeared, participants were required to cancel the *release* of the button on the side corresponding to the change in circle colour (that is, they had to maintain force on that side, and refrain from releasing the button) but to continue with the release of the button on the other side. If the participant did this successfully the trial was counted as successful, and feedback appeared in the feedback window reading “correct”. All other responses (release of both buttons, release of the wrong button, release of no buttons) resulted in a trial being deemed incorrect, and the feedback message displayed “incorrect”. Adjustments to SSD were performed in the same way as those in the press condition described above, based on trial success.

The release-ignore condition featured 300 trials, divided into 200 go trials, 50 stop trials and 50 ignore trials. The ignore trials operated in a similar manner to those in the press-ignore condition, though the behavioural imperative associated with them differed. As with go and stop trials in this condition, these trials started with a screen informing the participant to press and hold both buttons. Following the fixation cross, the go stimulus appeared. After SSD, the ignore signal occurred on either the left or right side, though this needed to be ignored (that is, a bimanual button release was still required). The proportion of trials involving additional signals (i.e., stop or ignore signals) in the press-ignore and release-ignore conditions (1/6 of trials were ignore; 1/6 of trials were stop) was consistent with the proportion of additional stimuli in conditions *without* ignore trials (in which 1/3 of trials were stop trials). Warnings reading “you’ve slowed down” also occurred in both release and release-ignore conditions, though the average RTs from the release-go-only block were used as a reference, rather than the press-go-only block.

**Figure 2:**
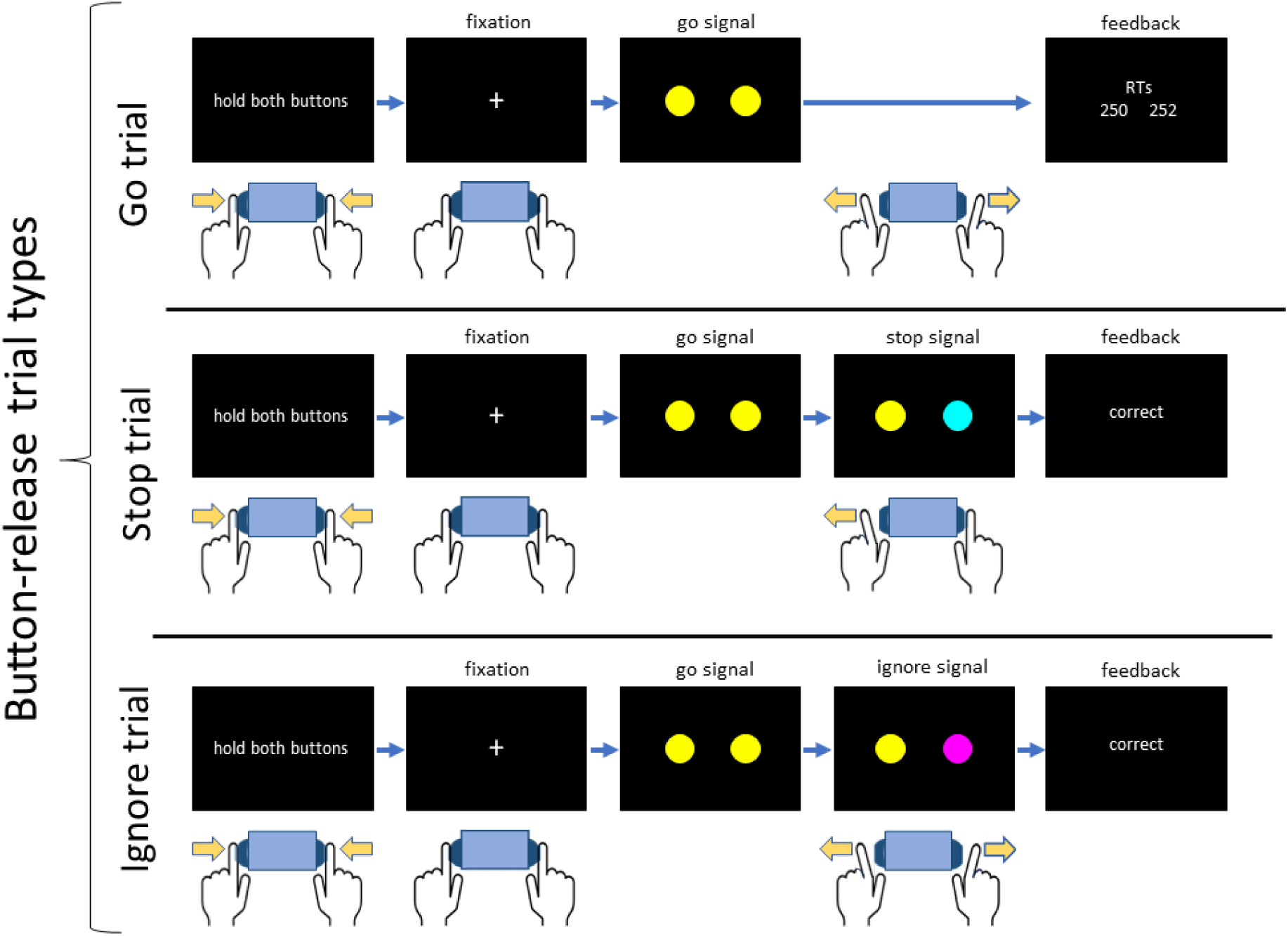
Trial types occurring in the two conditions requiring button releases as the primary response. Note that the “release” condition featured only go and stop trials, while the “release-ignore” condition also included ignore trials. Trials began with text informing participants to hold both buttons. Once both buttons were depressed, this triggered the transition to the fixation cross. The imperative associated with the go signal was to release both buttons. The stop signal (here, a cyan circle) could appear on either the left or right side and occurred after a variable stop signal delay (SSD, described in main text). Participants were required to refrain from releasing the button on the side of the stop signal. Ignore signals (occurring in ignore trials, in the third row), also occurred after SSD, but required a bimanual button release, as in go trials.

### 2.8 Data Processing and Analysis

Statistical analyses were conducted using linear mixed models (LMMs) and generalised linear mixed models (GLMMs) run within the statistical package Jamovi (The Jamovi Project, 2021) which operates via the statistical language R (R Core Team, 2021). GLMMs were run using the Jamovi module GAMLj (Gallucci, 2019). Model selection (i.e., determination of the random effects structure to be used) was performed based on BIC (Schwarz, 1978).

#### 2.8.1 Go Trial Reaction Time Analysis

A GLMM was run on RT values (averaged across left and right hands) from correct go trials with condition as a sole fixed factor with 6 levels (Go-Only-Press, Go-Only-Release, Press, Release, Press-Ignore, Release-Ignore). When anticipating the possible need to inhibit an upcoming movement, adjustments may be made within inhibitory networks to facilitate the potential upcoming action cancellation (Lavallee et al., 2014). These changes are referred to as proactive inhibition. Behaviourally, a manifestation of these changes is proactive slowing, which is quantified by comparing a participant’s reaction time (RT) in go trials during stopping tasks to RTs during a block without stop signals (van de Laar et al., 2011). Here, we investigated this in each condition using Bonferroni corrected post-hoc tests, comparing the go responses from the stop signal tasks to the go responses from the go-only blocks which used the same movement type (i.e., press or release).

#### 2.8.2 Analysis of Stopping Delays and Ignore Delay

Stopping delays or, the “stopping interference effect” refer to the delays in the continued response during response-selective stopping relative to reaction times in go trials (Gronau et al., 2024; Wadsley et al., 2022) A trial level measure of stopping delays was determined by subtracting the mean go trial reaction time from the RT of the non-stopped hand in stop trials (Wadsley et al., 2023b). To avoid complications regarding missing levels (due to no ignore trials in the A LMM (Gaussian distribution and identity link) was run only on the data from the stimulus selective conditions using go movement (press, release) and trial type (stop, ignore) as fixed factors. The ultimate random effects structure was a maximal model including participant intercepts, slopes for both factors, and the interaction term.

#### 2.8.3. Processing of EMG data

EMG data processing was performed using MATLAB (MathWorks, 2018). EMG signals were digitally filtered using a fourth-order band-pass Butterworth filter at 20–500 Hz. In the press and press-ignore conditions, the precise onset and offset times of EMG bursts were detected using a single-threshold algorithm (Hodges & Bui, 1996): Signals from each trial were rectified and low-pass filtered at 50 Hz. Following this, we used a sliding window of 500ms to find the trial segment with the lowest root mean squared (RMS) amplitude, which was used as a baseline. EMG bursts were identified when the amplitude of the smoothed EMG was more than 3 SD above baseline. EMG bursts separated by less than 20ms were merged together to represent a single burst. Following identification of the onset and offset times, we used time constraints to identify two types of EMG burst: The *RT-generating burst* was identified as the last burst with an onset occurring *after* the go signal and *prior* to the recorded button press. Some stop and ignore trials also exhibited *partial bursts*, i.e., EMG bursts that started to decrease in amplitude before enough force was generated to trigger an overt button press. Partial bursts were identified in each hand as the earliest burst where peak EMG occurred *after* the presentation of a stop/ignore signal but *prior* to the onset of the RT-generating burst from the responding hand. Furthermore, peak EMG amplitude of a partial burst was required to be greater than 10% of the average peak amplitude from that participant’s successful go trials in that condition. Critically, these time and amplitude constraints exclude activity occurring *after* the RT-generating burst, as this is likely to represent mirror activity or activity unrelated to the task. Partial bursts which were separated by less than 20ms were merged into a single burst, to further reduce the possibility of mirror activity being classified as a partial burst. For each burst (RT-generating and partial), we extracted the time of burst onset and offset, as well as the time of peak amplitude.

In the release release-ignore conditions, two distinct types of EMG landmark were identified (analogous to the “RT generating” and “partial” bursts identified in the press and press-ignore conditions). For the RT generating burst, the point at which EMG amplitude became <10% of the mean amplitude measured from the start of EMG recording (400ms before the go signal) to the go signal, was determined as EMG offset time. If the algorithm failed to detect a drop to 10% of mean amplitude between the go signal and the recorded behavioural response, the threshold was incremented by one percent (e.g., from 10% to 9%), and the process was repeated. The other type of response was a “partial release” which were identified only on stop and ignore trials. These were identified between SSD and the EMG offset time in the responding hand when the EMG amplitude (in the stopping hand) became less than 50% of that in the pre-trial phase. Following this, it was also required that EMG amplitude returned to the amplitude of the hold phase.

EMG envelopes were obtained using full-wave rectification and a low-pass filter at 10 Hz. For each subject, the amplitude of EMG envelopes from each hand was normalised by the average peak EMG from successful bimanual go trials in the press condition, thus allowing direct comparisons between participants and conditions.

#### 2.8.3. CancelTime

In successful stop trials and ignore trials in the press and press-ignore conditions featuring a partial burst, the latency of the peak amplitude of the partial EMG burst, relative to that trial’s SSD, can be used to determine a trial-level estimate of stopping latency, referred to as CancelTime (Jana et al., 2020; Raud et al., 2022). While CancelTime is conventionally calculated based on partial burst latency in the stopping hand, we expand upon this definition here, calculating CancelTime based on partial bursts in either hand. To identify an equivalent measure in the release conditions, we calculated the time between SSD and the lowest point of the partial release (see Figure 3 and section 2.8.2). Thus, in both conditions CancelTime represents the point at which muscle activity demonstrates a reactive change to motor plans in response to the stop signal. In the press conditions this involves a suppression of muscle activity (a reduction in amplitude), and in the release conditions this involves an *increase* in muscle activity (an increase in amplitude).

**Figure 3:**
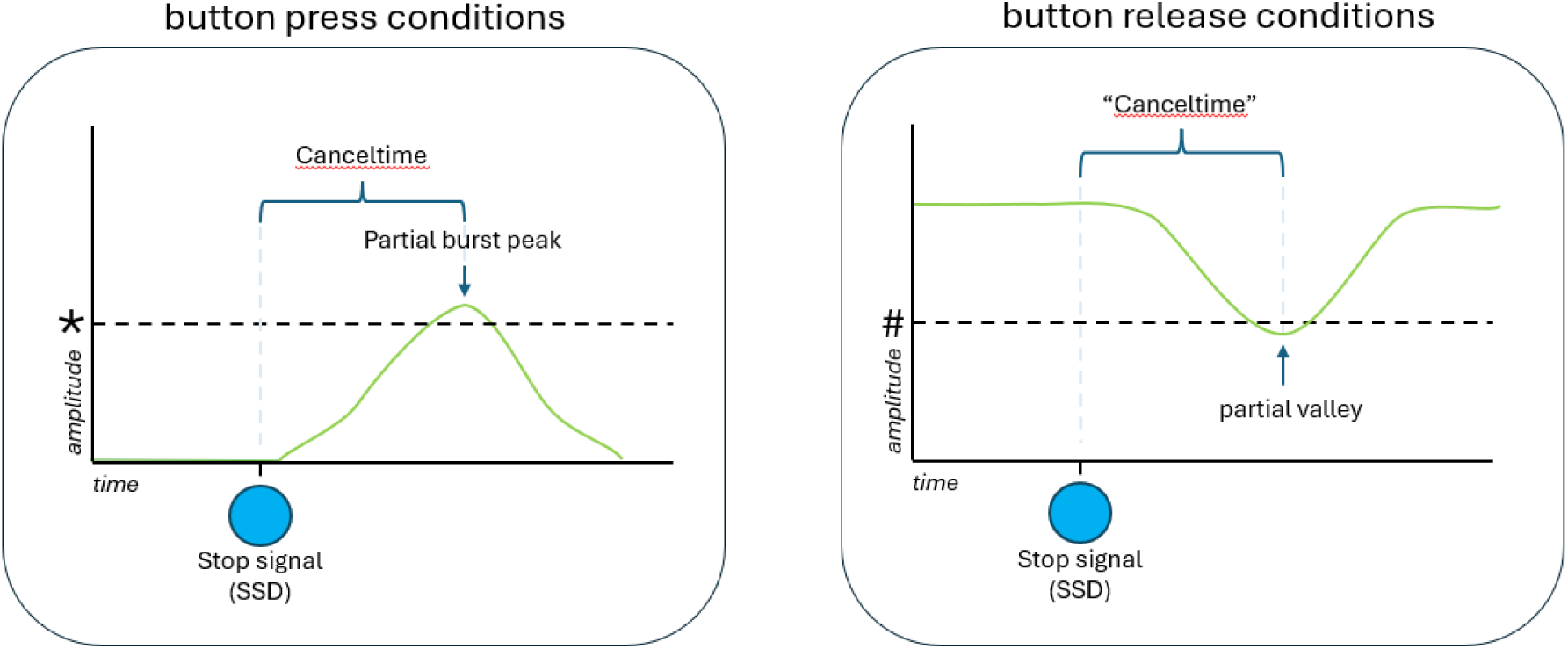
The green line represents EMG amplitude during successful stop trials in which the go response was initiated, but did not result in an overt behavioural response (i.e., a button press or button release). CancelTime is calculated as the time between the presentation of the stop signal (SSD) and the time at which cancellation of the go response begins (the peak of the partial burst in press conditions, and the lowest point of the valley in release conditions). For detail regarding the categorisation of these landmarks refer to section 2.8.2. **\*** Detection of partial bursts required that amplitude exceed 10% of that participant’s mean go trial amplitude in that condition. **#** Detection of partial releases required a release to <= 50% of mean pre-trial hold-phase amplitude for that trial.

For trials featuring partial bursts in both hands, we used the latency of whichever partial burst occurred earlier. To minimise the influence of outliers, any CancelTimes that were greater than1.5 x the interquartile range above the third quartile (performed on a per-condition and per-trial-type basis) were excluded (Jana et al, 2020). This removed 5.22% of stop trials and 6.6% of ignore trials with partial responses. CanceTimes that were shorter than 50ms, were also removed, as these likely reflect anticipatory responses rather than reactions to the stop signal (7.23% of stop trials and 8.48% of ignore trials with partial bursts/releases).

An initial preliminary GLMM revealed no difference in CancelTime between stop trials between conditions with and without ignore stimuli *χ*^2^(1) = 0.19, *p* = 0.662. Accordingly, CancelTime data were combined across conditions with/without ignore trials to avoid complications relating to missing levels for the trial type variable (stop vs ignore) in the press and release conditions which did not feature ignore trials.

Following this, a generalized linear mixed model (GLMM) with a gamma distribution and a log link function was run on this trial-level data. Fixed factors included go-movement (press/release), trial type (stop/ignore), and partial-presence (whether partial bursts/releases were observed bilaterally or unilaterally). Any trials which featured bimanual partial responses, that were not synchronous (a difference in latency of >50ms), were categorised as unimanual, and the latency of the earlier burst was taken. This rule was included to ensure that the partial responses deemed to occur bimanually plausibly reflected a unitary inhibitory response, rather than two distinct inhibitory reactions. In the press conditions 63.66% of partial bursts in stop and 50.31% of partial bursts in ignore trials were synchronous. In the release condition 32.3% of ignore trials and 37.37% of stop trials. The final random effects structure, following model section based on BIC, included participant intercepts and slopes for partial-presence.

## 3. Results

### 3.1 Accuracy

Mean accuracy in go trials was over 98% for all conditions and accuracy in ignore trials was >99% in both conditions featuring ignore trials. These results suggest high levels of attention during the task. Accuracy in stop trials was 50.10% in the press condition, 45.09% in the release condition, 48.94% in the press-ignore condition, and 44.51% in the release-ignore condition, indicating that the staircase algorithm on SSD was effective.

### 3.2 Go trial reaction times

Mean (and 95% CI) RTs for correct responses in each trial type are presented in Table 1 and RT distributions for each condition are represented in Figure 3. The go trial reaction time analysis revealed a significant main effect of condition, *χ^2^* (5) = 97.09, *p* < 0.001. Bonferroni-corrected post-hoc tests revealed significant proactive slowing in the press condition (*z* = 9.31, *p* < 0.001) and press-ignore condition (*z* = 7.67, *p* < 0.001) relative to the press go-only condition. Significant proactive slowing was also observed in the release condition (*z* = 7.09, *p* < 0.001) and release-ignore condition (*z* = 5.86, *p* < 0.001) relative to the release go-only condition. Furthermore, RTs in the release condition were significantly slower than those in the release-ignore condition (*z* = 3.28, *p* < 0.016), the press condition (*z* = 3.15, *p* < 0.024), and the press-ignore condition (*z* = 4.27, *p* < 0.001). No other comparisons between the main experimental conditions were statistically significant.

**Table 1:** Reaction Times (RT, ms) from each trial type. 95% confidence intervals in parentheses.

| Condition | Trial | Correct RT (ms) | Failed RT(ms) |
| --- | --- | --- | --- |
| **Press go-only** | Go | 292 (77) | * |
| **Press** | Go | 422 (105) | * |
|  | Stop | 467 (129) | 381(87) |
| **Press-ignore** | Go | 416 (99) | * |
|  | Stop | 488(149) | 390(95) |
|  | Ignore | 485(159) | * |
| **Release go-only** | Go | 318 (118) | * |
| **Release** | Go | 481 (141) | * |
|  | Stop | 565(185) | 488(186) |
| **Release-ignore** | Go | 435 (110) | * |
|  | Stop | 535(184) | 438(175) |
|  | Ignore | 505(163) | * |

\*Given the very small number of failed go and ignore trials (<98%) means and SDs are not reported

**Figure 4:**
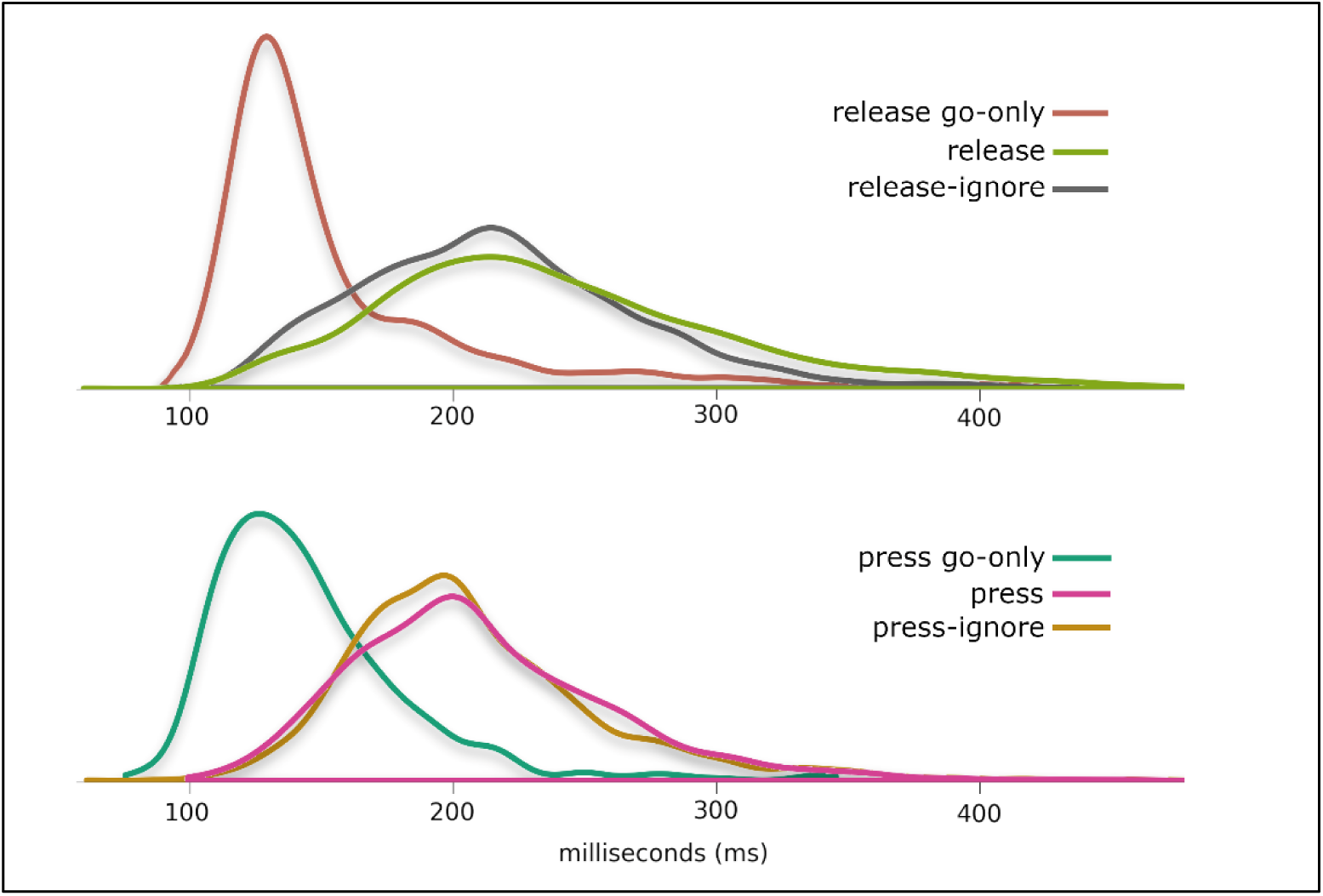
Density plots representing the distributions of reaction times (RT) for go trials in each condition. Top plot shows release conditions, and bottom plot shows press conditions

### 3.3 Stopping delays and ignore delays

The results of the model run on stopping delays are depicted in Figure 5. The model revealed no significant main effect of movement *F*(1,19) = 1.60, *p* = 0.220, a significant main effect of trial type, *F*(1,19) = 128.08, *p* < 0.001, and a significant interaction effect between these two variables *F*(1,19) = 5.58, *p* = 0.029. Tests of simple main effects revealed that, when comparing press and release conditions there was a greater difference between stop trials (*F*[1,19] = 3.52, *p* = 0.076) than ignore trials (*F*[1,19] = 0.02, *p* = 0.886) though neither difference was statistically significant. A post-hoc comparison run on trial-type revealed stopping delays in response to stop trials (191ms) were significantly greater than those for ignore trials (71ms), *t*(19) = 11.32, *p* < 0.001. Estimated marginal means for stopping delays in stop trials were 169ms [95%CI = 123ms-216ms] and 212ms [95%CI = 177ms – 247ms] in the press and release conditions, respectively. Estimated marginal means for stopping delays in ignore trials were 70ms [95%CI = 32ms-107ms] and 72ms [95%CI = 30ms – 114ms] in the press and release conditions, respectively.

**Figure 5:**
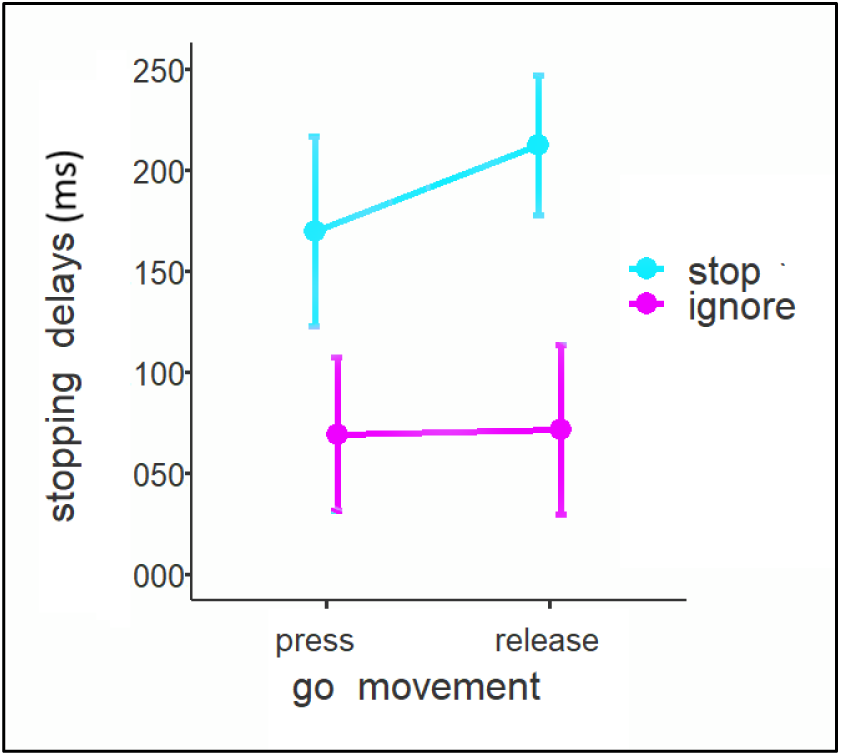
Stopping delays (also referred to as the stopping interference effect) following stop and ignore signals in the Press-ignore and Release-ignore conditions. Error bars represent 95%Cis.

### 3.4 CancelTime

Figure 6 depicts average (group level) EMG profiles from stop and ignore trials in press and release conditions. Partial bursts are observable in the press conditions as transient increases in muscle activity followed by suppression, and “partial release” in the release conditions are observable as a decrease in EMG followed by an increase back to (and beyond) the baseline tonic level.

**Figure 6:**
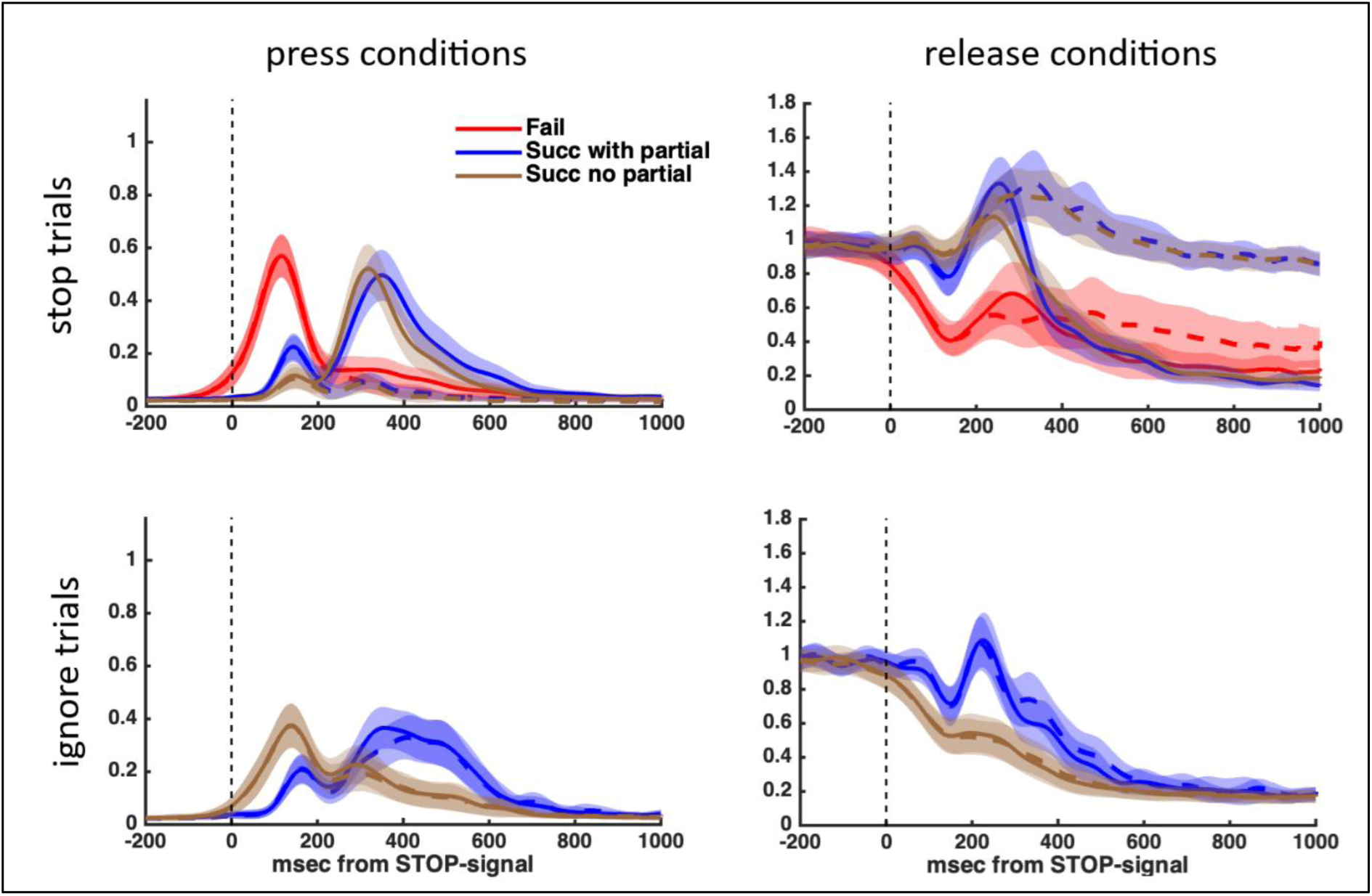
Average (group) EMG profiles representing the mean EMG amplitude over time, synchronised to the presentation of the stop or ignore signal (vertical dashed lines). Solid lines represent the hand on the same side as the stop/ignore signal, while dashed lines represent the other (non-cued) hand. Partial bursts are observable prior a release of force in press conditions, while partial releases are observable prior to an increase in force in the release conditions. Notably, both types of partial response (presses and releases) are observable in both hands (dashed and solid blue lines).

The model run on EMG-derived CancelTime revealed a significant higher order interaction effect between go-movement (press/release), trial-type (press/ignore) and partial-presence (bilateral/unilateral) *χ*^2^(1) = 6.49, *p* < 0.011., which was investigated with a test of simple main effects of go movement split by trial type and partial-presence. The analysis revealed a significant difference in CancelTime between press and release conditions in stop trials where stopping was only observed in one hand *χ*^2^(1) = 75.70, *p* < 0.001), whereby CancelTime in release conditions (178ms) was slower in press conditions (130ms). However, when partial responses (bursts or releases) occurred bilaterally there was no significant difference in CancelTimes between press (143ms) and release (144ms) conditions *χ*^2^(1) = 0.03, *p* = 0.863. There was also no significant difference in CancelTimes in response to ignore trials when stopping occurred bilaterally (press = 169ms: release = 156ms; *χ*^2^[1] = 2.06, *p* = 0.152), nor when it was observed unilaterally (press = 173ms: release = 173ms; *χ*^2^[1] = 0.00, *p* = 0.972).

The model also revealed a significant interaction between go movement and trial type *χ*^2^(1) = 20.59, *p* < 0.01. This was driven by there being a significant effect of go movement for stop trials (release = 160ms: press = 136ms: *χ*^2^[1] = 39.12, *p* < 0.001), but not for ignore trials (release = 164ms: press = 170ms: *χ*^2^[1] = 1.18, *p* = 0.277). There was also a significant interaction between go movement and partial-presence *χ*^2^(1) = 18.46, *p* < 0.001. When averaging across trial types, there was no significant effect of go movement in trials with synchronous partial bursts (release = 150ms: press = 155ms: *χ*^2^[1] = 1.18, *p* = 0.277), though there was a significant effect of go movement when partial bursts were unilateral, or asynchronous (release = 175ms: press = 150ms: *χ*^2^[1] = 28.66, *p* < 0.001). There was no significant interaction between trial type and partial-presence *χ*^2^(1) = 0.02, *p* < 0.884.

The model also revealed a significant main effect of go-movement *χ*^2^(1) = 7.01, *p* = 0.008, whereby overall, CancelTime was faster in the press conditions (152ms) than the release conditions (162ms). There was also a main effect of trial type *χ*^2^(1) = 32.09, *p* < 0.001, whereby overall CancelTime was faster to stop trials (147ms) than ignore trials (167ms). No significant main effect of partial-presence was observed *χ*^2^(1) = 1.81, *p* = 0.179.

A follow-up Bayesian t-test using the default Cauchy prior distribution with a scale factor of 0.707 (Wagenmakers et al., 2018) was conducted on the trials with synchronous bimanual partial responses to determine whether the lack of a significant difference between press and release conditions reflected a true null result. The result revealed moderate evidence for the null BF01 = 8.49.

**Figure 7:**
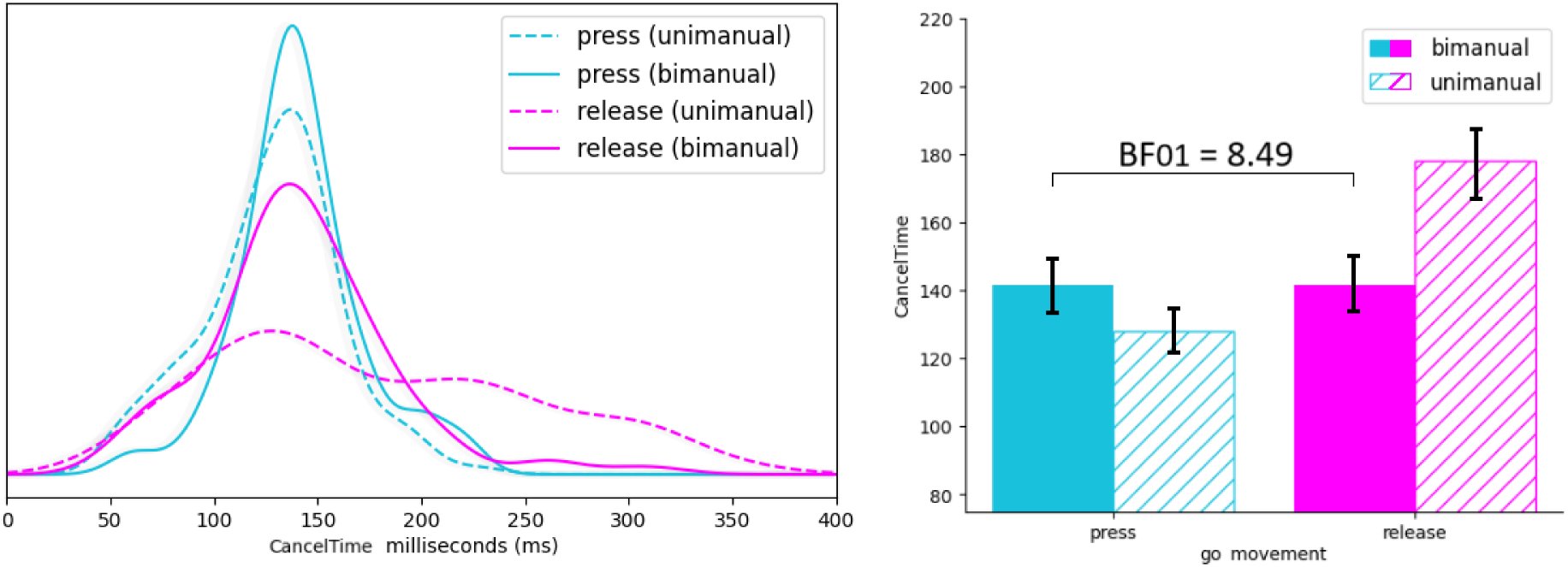
The left panel depicts CancelTime distributions (ms) in stop trials from press and release conditions split by whether there was synchronous physiological evidence of non-selective action cancellation (partial responses observed with electromyography in both hands), or whether this was observed only in one hand.

## 4. Discussion

The current study used novel behavioural paradigms and electromyography to assess the latency of reactive action cancellation in contexts requiring either muscle suppression or muscle contraction. No difference in stopping latencies between these contexts was observed when physiological evidence of synchronous action cancellation in both hands was present. The observation that cancelling a voluntary response via an increase in muscle activation can be performed as quickly as when this is achieved via muscle suppression raises fundamental questions regarding the specialised nature of inhibitory mechanisms, and how these manifest at the level of the muscle.

Past research has described EMG-derived CancelTime (the time from presentation of a stop signal until suppression of an initiated movement begins) as inhibitory signals from specialised inhibitory networks arriving at the level of the muscle (Jana et al., 2020). Furthermore, some interpretations have associated CancelTime with specific components of theoretical models of the stopping network. For example, by one account, action cancellation is underpinned by distinct “pause” and “cancel” mechanisms. The pause mechanism purportedly occurs broadly in response to attentional capture, is faster than the cancel response, and is non-selective (that is, it effects all ongoing actions, rather than a subset). In contrast, the selectively applied cancellation of an action (i.e., to one effector and not others) can only be mediated by a distinct, slower, and consciously controlled response (referred to as a “cancel”; Diesburg & Wessel, 2021; Schmidt & Berke, 2017). EMG-derived CancelTime has previously been associated with the pause mechanism in particular, based on the rapid latency with which it occurs (Tatz et al., 2021), and the fact that attentional capture (even in contexts where stopping is not required) is sufficient to trigger it (Weber et al, 2023). However, the current result implies either that the inhibitory pause process must also accommodate muscle contraction, or that another response - distinct from the pause process, but just as rapid - can also occur upon presentation of the stop signal.

The notion that partial releases in the release condition may have been a result of a pause process would require that this mechanism manifests differently depending on contextual demands. That is, advance knowledge may allow this same automatic mechanism to trigger an *increase* in muscle activation to reinstate the tonic contraction (i.e., keep the button pressed). While the neurological pathway proposed to underpin this process (a network originating in the inferior frontal gyrus and involving the sub-thalamic nucleus and globus pallidus interna; Chen et al., 2020) is typically associated with an inhibitory function, a number of questions remain regarding its precise function (Bingham et al., 2023), and the possible role of other connections. As such, the notion of a generalised reactive mechanism that relies on this network is plausible. Furthermore, the fact that the absence of a difference in CancelTime between press and release conditions was only observable when there was evidence of action cancellation in both hands (i.e., when it was demonstrated that the stop process was not deployed selectively) lends support to this interpretation - given that the pause process is purportedly applied globally, rather than to a subset of effectors (Diesburg & Wessel, 2021). Nonetheless, recent research using near-infrared spectroscopy observed that the presence of a partial burst was not associated with increased activity in the inferior frontal gyrus, as would be expected if the pause process was the primary mediator of the inhibition observed in partial bursts (Kemp et al., 2024). Given the number of unmeasured mediators occurring between the activation of these cortical regions and the activation/suppression of muscle fibres, it remains that the presence of partial bursts represents an ambiguous combination of neurological mechanisms. While partial bursts represent trials in which it is known that inhibition occurred, as there is direct empirical evidence of this, the lack of neurological evidence in brain imaging contrasts may result from the fact that the point of contrast (trials *without* partial bursts) likely reflects a combination of trials with and without inhibition. That is, inhibition may have occurred prior to the point at which muscle activation in response to the go signal (which must be present for the suppression of movement to be observable) begins. Nonetheless, future brain imaging and brain stimulation techniques may shed further light on the neural underpinnings associated with partial activations.

Another explanation is that that the partial releases observed in the release conditions are unrelated to the pause process and instead represent a distinct reactive bimanual go response that occurs with a similar latency to the stop process. Notably, this explanation raises questions regarding the functional utility of a specialised inhibitory system: If the initiation of an oppositional action can be performed just as quickly as typical action cancellation, to what extent is the cancellation of movement actually reliant on the suppression of muscle activity? The current result suggests that focussing solely on the suppression of muscle activity in inhibitory control research may over-emphasise the generalisability of this process to real world stopping contexts, in which stopping can be performed just as quickly via the initiation of a new, opposing movement. In many contexts, such as the scenario described where a reaching movement is cancelled upon observation of an insect, the facilitation of opposing muscle activity may be the primary mechanism by which the behavioural response occurs. On the other hand, because most action cancellation requires a combination of both suppression and facilitation of different muscle groups, it is perhaps unsurprising that both reactive movement initiation and reactive cancellation can both occur in a similarly rapid fashion, and the possibility of a specialised mechanism to mediate both of these processes is plausible. Future research could collect EMG data from agonist/antagonist muscle pairs in similar tasks to those used in the current study, to investigate these effects in opposing muscle groups.

In sum, the current findings indicate that cancelling a movement via the reinitiation of tonic force can be performed just as rapidly as when this is achieved via the suppression of muscle activity. This challenges assumptions relating to the pause then cancel model (specifically, that the pause function is purely inhibitory) and suggests the existence of highly generalised reactive action updating mechanisms. Future work, combining brain imaging techniques with similar action cancellation tasks could serve to better characterise potential dissociations between distinct stopping mechanisms.

## Footnotes

1 This variable pre-cue period served to reduce the predictability of the subsequent go signal onset and ensure that participants were responding to the visual stimulus, rather than predicting its appearance (Verbruggen et al., 2019).

